# Enzyme-based synthesis of single molecule RNA FISH probes

**DOI:** 10.1101/248245

**Authors:** Christian Lanctôt

## Abstract

Single molecule RNA fluorescence *in situ* hybridization (smRNA FISH) allows the quantitative analysis of gene expression in single cells. The technique relies on the use of pools of end-labeled fluorescent oligonucleotides to detect specific cellular RNA sequences. These fluorescent probes are currently chemically synthesized. Here I describe a novel technique based on the use of routine molecular biology enzymes to generate smRNA FISH probes without the need for chemical synthesis of pools of oligonucleotides. The protocol comprises 3 main steps: purification of phagemid-derived single stranded DNA molecules comprising a segment complementary to a target RNA sequence; fragmentation of these molecules by limited DNase I digestion; and end-labeling of the resulting oligonucleotides with terminal deoxynucleotide transferase and fluorescent dideoxynucleotides. smRNA FISH probes that are obtained using the technique presented here are shown to perform as well as conventional probes. The main advantages of the method are the low cost of probes and the flexibility it affords in the choice of labels. Enzyme-based synthesis of probes should further increase the popularity of smRNA FISH as a tool to investigate gene expression at the cellular or subcellular level.

## INTRODUCTION

The most widely used methods to determine gene activity, such as qRT-PCR, RNA-seq or microarray analysis, provide expression levels averaged over entire cell populations. In recent years, the growing use of methods designed to measure gene expression in single cells has revealed features that cannot be deduced from averaged data, i.e. the pulsatile nature of transcription and the large cell-to-cell variability in expression in isogenic populations [1]. One of these methods is single molecule RNA FISH (smRNA FISH), which was pioneered by the Singer group [2, 3] and further developed by others (for review see [4]). This technique was used to gain important insights on cell-to-cell variability in stem cell niches, cellular differentiation and signaling, among others [5-7]. smRNA FISH can be used to analyze gene expression in a variety of models, including cultured cells, tissue sections and whole embryos [8-11]. Like most imaging-based techniques, the main drawbacks of smRNA FISH are its low throughput and its inability to interrogate whole genomes. However, advances in automated image acquisition and analysis coupled with error-free combinatorial labeling promise to overcome these limitations, as witnessed by the recent development of a multiplex smRNA FISH protocol (MERFISH) that enabled the imaging of hundreds of distinct RNA species in thousands of individual cells [12, 13].

In its present form, the smRNA FISH technique uses single stranded end-labeled oligonucleotides as probes [14]. These oligonucleotides are usually 20-30 nucleotides in length and harbor a single fluorophore covalently linked to their 3’ end. A typical probe comprises a pool of 24-48 such fluorescent oligonucleotides, each hybridizing to a different sequence on the target RNA. An alternative protocol (smiFISH) has recently been developed in which the oligonucleotides complementary to the target RNA are not themselves fluorescent; rather, all of them contain an identical tail of ≥28 nucleotides which serves as the target for a unique fluorescent oligonucleotide, referred to as the FLAP probe [15]. In both cases, since the composition of the probe is known and hence so is the expected number of fluorophores per target molecule, the intensity of the signal is similar for each hybridized RNA. Furthermore, since individual RNA molecule are smaller than the optical resolution of the light microscope, the fluorescent signals appear as diffraction-limited spots. These properties allow the assignment of spots of similar intensity and ‘shape’ to single RNA molecules [16]. In yet another embodiment of the smRNA FISH technique (bDNA-sm-FISH), the hybridization signal is amplified via the formation of a branched DNA complex which harbors multiple sites for the binding of the detection probe [17].

At present, probes for smRNA FISH are chemically synthesized. The procedure relies on well-known methods of solid-phase synthesis of nucleic acids. Probes are commercially available, yet their high cost can be prohibitive. Furthermore, once it has been synthesized, the pool of labeled oligonucleotides always harbor the same fluorescent dye, thereby limiting the flexibility of the technique. Here I present a novel and simple enzyme-based approach to the synthesis of smRNA FISH probes, which is both affordable and flexible and which should help to further promote the use of this powerful technique for the analysis of gene expression.

## MATERIALS AND METHODS

### Enzymes and reagents

Restriction enzymes were purchased from New England Biolabs (www.neb.com) or from ThermoFisher Scientific (www.thermofisher.com). Terminal deoxynucleotidyl transferase (Fermentas brand) was purchased from ThermoFisher Scientific. DNase I (2 U/μl) and dye-labeled dideoxynucleotide triphosphates were purchased from Jena BioScience (www.jenabioscience.com). Synthetic oligonucleotides were obtained from Sigma Aldrich (www.sigma-aldrich.com). Cell culture reagents were from ThermoFisher Scientific unless otherwise stated. Chemicals were obtained from Sigma Aldrich unless otherwise stated. The plasmids pGEM-3zf(−) and pGEM-3zf(+) were purchased from Promega (www.promega.com).

### Cell culture

HepG2 cells (ATCC no. HB-8605, obtained from another laboratory) were cultivated in Dulbecco’s modified essential medium (DMEM) supplemented with 10% (v/v) fetal bovine serum, 100 U/ml penicillin, 100 μg/ml of streptomycin and 1X non-essential amino acids. When confluent, cells were passaged by a short (~4 minutes) treatment with 0.25% trypsin/0.91mM Na-EDTA and seeded at 20,000 cells/cm^2^.

### RNA extraction and synthesis of first strand cDNA

Total RNA was extracted and purified from confluent HepG2 cells using the RNEasy kit according to the manufacturer’s instructions (Qiagen, www.qiagen.com). Five micrograms of total RNA were reverse-transcribed into first strand complementary DNA in the presence of 0.5 μg/ml of dT_18_ primer (ThermoFisher Scientific) and 10 U/μl of the Superscript III enzyme (ThermoFisher Scientific). Incubation was performed at 42°C for 10 minutes and 50°C for 60 minutes, after which the reverse transcriptase was inactivated by incubation at 70°C for 15 minutes. The first strand cDNA was used in downstream PCR reactions without further purification.

### Cloning of target RNA sequences in phagemid vector

DNA fragments corresponding to target RNA sequences were obtained by PCR. The sequences of forward and reverse primers that were used to amplify part of the human CREB3, TLN1, SENP3 and POLR2A sequences are given in Supplementary Table 1. After purification, the blunt DNA fragments (~80-120 ng) were ligated into 50 ng of SmaI-linearized pGEM-3zf(−) phagemid. The ligation reaction was transformed into competent *E. coli* cells of the F’-containing JM109 strain (Promega). Transformants were grown on plates containing IPTG (isopropyl-β-D-thiogalactoside) and X-Gal (5-bromo-4-chloro-3-indolyl β-D-galactopyranoside), which allowed blue/white selection of vectors with inserts. Positive clones were validated by sequencing.

### Growth and purification of ssDNA molecules

Vectors which contain the insert in the orientation opposite to that of the f1 origin are used to generate antisense probes. Vectors in which the insert and the f1 origin are in the same orientation are used to generate control sense probes, if needed. Single stranded DNA (ssDNA) molecules were isolated using a protocol based on published methods [18, 19]. Briefly, clones were grown in 1 ml of LB medium supplemented with ampicillin (75 μg/ml) for 14-16 hours at 37°C with agitation. This culture was diluted 1:100 (v/v) in 40 ml of the same medium and further grown at 37°C until the absorbance of the culture at 600 nm reached ~0.08. At that time, a 80 μl aliquot of M13KO7 helper phage (New England Biolabs) was added, for a final concentration of 2 × 10^8^ pfu/ml (multiplicity of infection ~ 3). Infection was allowed to proceed for 1.5 hour at 37°C with agitation. Kanamycin sulfate was then added at a final concentration of 70 μg/ml to select for cells infected with the helper phage. The temperature was lowered to 30°C and cells were further cultivated for 12-14 hours to allow for production and release of ssDNA.

The bacterial culture was centrifuged at 11,000 *g* for 10 minutes at 4°C. Thirty-five ml of supernatant were transferred into a new 50 ml tube, taking care not to disturb the bacterial pellet. Seven ml of 25% polyethylene glycol-8000/2.5 M NaCl was added to the supernatant. After thorough mixing, the solution was incubated on ice for 2-4 hours. The precipitated phage particles were centrifuged at 11,000 *g* for 10 minutes at 4°C. The supernatant was decanted and drained completely. The pellet was resuspended in 1 ml of 1X TE (10 mM Tris-HCl pH 8/1 mM EDTA) and transferred to a 2 ml microtube. An equal volume of chloroform was added. The mixture was thoroughly vortexed (at least 30 seconds at high speed) and centrifuged at 11,000 *g* for 5 minutes at 4°C. Typically, a white pastille of polyethylene glycol was found at the interface. The top aqueous phase was recuperated and transferred to a new microtube. Nucleic acids were then extracted by phenol/chloroform following standard protocols. Typically, a total of 0.7 ml was recuperated after organic extraction. Nucleic acids were ethanol-precipitated and resuspended in 50 μl of 5 mM Tris-Cl pH 8.5. The concentration of ssDNA was determined by measurement of absorbance at 260 nm using a Nanodrop instrument. The integrity and purity of the ssDNA were ascertained by agarose gel electrophoresis. The typical yield from a 40-ml culture was 20-30 μg of ssDNA. Before proceeding to the next step, an aliquot of 10 μg was further purified on a Zymo Research DNA Clean-5 column (www.zymoresearch.com). Elution was performed with 10 μl.

### Generation of short oligonucleotides from single stranded DNA molecules

Three reactions were assembled on ice in 0.2 ml thin-walled PCR tubes. Each contained one microgram of ssDNA and was completed to 7 μl with distilled water. To each reaction was added 2 μl of 5X DNase I reaction buffer (final concentrations: 10 mM Tris-Cl pH 8/2.5 mM MgCl_2_/0.5 mM CaCl_2_). One microliter of water was added to the first PCR tube to serve as a non-digested control. Two dilutions of DNase I were prepared in water: 1:20 (v/v, final concentration of 0.1 Kunitz U/ml) and 1:40 (v/v, final concentration of 0.05 Kunitz U/ml). Optimal dilutions must be determined empirically (see Results). Dilutions were kept on ice. One microliter of a given dilution was added to a different PCR tube. The PCR tubes were immediately transferred to a pre-heated PCR cycle programmed for an incubation of 15 minutes at 65°C followed by an incubation of 3 minutes at 95°C.

After DNase I digestion, the size of the DNA products was analyzed on a 12% denaturing polyacrylamide vertical gel electrophoresis. Aliquots of 1 μl of each DNase I reaction and of the control reaction (corresponding to 100 ng of ssDNA) were run on gel. A molecular marker consisting of an equimolar mixture of single stranded oligonucleotides of known lengths (in the present case 20 nt, 30 nt, 42 nt, 65 nt and 83 nt) was run alongside the samples. After migration, the gel was stained with 1X GelStar and visualized according to the manufacturer’s instructions (www.lonza.com).

### End labeling of short oligonucleotides with fluorescent dideoxynucleotides

The DNase I reaction (9 μl) that contained fragments of approximately 20 to 60 nucleotides in length was chosen for labeling. Four microliters of distilled water, 4 μl of 5X TdT buffer (ThermoFisher Scientific), 2 μl of a 1 mM solution of fluorescent ddCTP (e.g. 5- propargylamino-ddCTP-Cy3) and 1 μl (20 U) of terminal deoxynucleotidyl transferase (TdT) were added directly to the chosen DNase I products. The labeling reaction was incubated for 60 minutes at 37°C. The TdT enzyme was then inactivated by incubation at 70°C for 10 minutes. The oligonucleotides were separated from the unincorporated dye-labeled ddCTP molecules by overnight ethanol precipitation. Labeled oligonucleotides were resuspended in 15 μl of Tris-Cl pH 8.5 and their concentration determined by measuring absorbance at 260 nm on a Nanodrop instrument. The absorption spectra of control and labeled products were determined on a Nanodrop instrument.

### Single molecule RNA FISH

The smRNA FISH protocol is based on previously published ones [8]. Cells were grown on gelatin-coated 18 mm × 18 mm #1.5 coverslips, fixed with 4% formaldehyde in 1X PBS for 10 minutes at room temperature, washed and stored overnight in 70% EtOH. After rehydration, hybridization was carried out in 2X SSC/25% formamide/10% dextran sulfate for 8-12 hours at 37°C. Each ssDNA-derived probe was used at a final concentration of 0.5-1 ng/μl. The Stellaris probe against human POLR2A was labeled with Quasar-670 and used at a concentration of 40 nM. The hybridization solution also contained 20 mM ribonucleoside vanadyl complex (New England Biolabs) and 200 μg/ml yeast tRNA to protect cellular RNAs. After washing with 25% formamide/2X SSC for 30 minutes at 37°C and twice with 2X SSC for 5 minutes at room temperature, samples were counterstained for 2 minutes at room temperature with 4’,6-diamidino-2-phenylindole (DAPI, 1 μg/ml in 2X SSC). After rinsing with 2X SSC, samples were mounted in Prolong Gold (ThermoFisher Scientific). Slides were cured at room temperature for 3 days before imaging.

### Image acquisition and analysis

Optical sections (25-35 at 250 nm intervals) were acquired on an Olympus IX71 inverted epifluorescence microscope using a 100X/numerical aperture 1.4 oil immersion objective and a Hamamatsu ORCA charged-coupled device (CCD) camera. The exposure time per optical section was 500-700 ms. The *xy* pixel size was 67 nm. The dynamic range of the images was 12 bits. Images were filtered using Gaussian kernels of 5 (for background subtraction) and 0.5 (for feature enhancement). The counting of cytoplasmic mRNAs, the measurement of their intensity and the construction of average mRNA molecules were performed using the freely available MATLAB-based FISH-QUANT program [16]. Colocalization analysis was performed using the ImageJ JACoP plugin [20]. Briefly, image stacks were filtered as above and manually-thresholded. The colocalization algorithm that was used was object-based, with a minimum particle size set at 15 pixels. Two objects were deemed to be colocalized if the distance between their geometrical centers was less than the optical resolution of the microscope setup. The colocalization value is given as the number of colocalized objects over the total number of objects.

## RESULTS

This article describes a new method to generate smRNA FISH probes using standard molecular biology techniques. The method comprises three steps, which are schematized on Figure 1 and described below. These are 1) the isolation of single stranded DNA molecules comprising a fragment complementary to the target RNA; 2) the fragmentation of these molecules into oligonucleotides; 3) the labeling of these oligonucleotides with a single fluorescent moiety.

**Figure 1.**
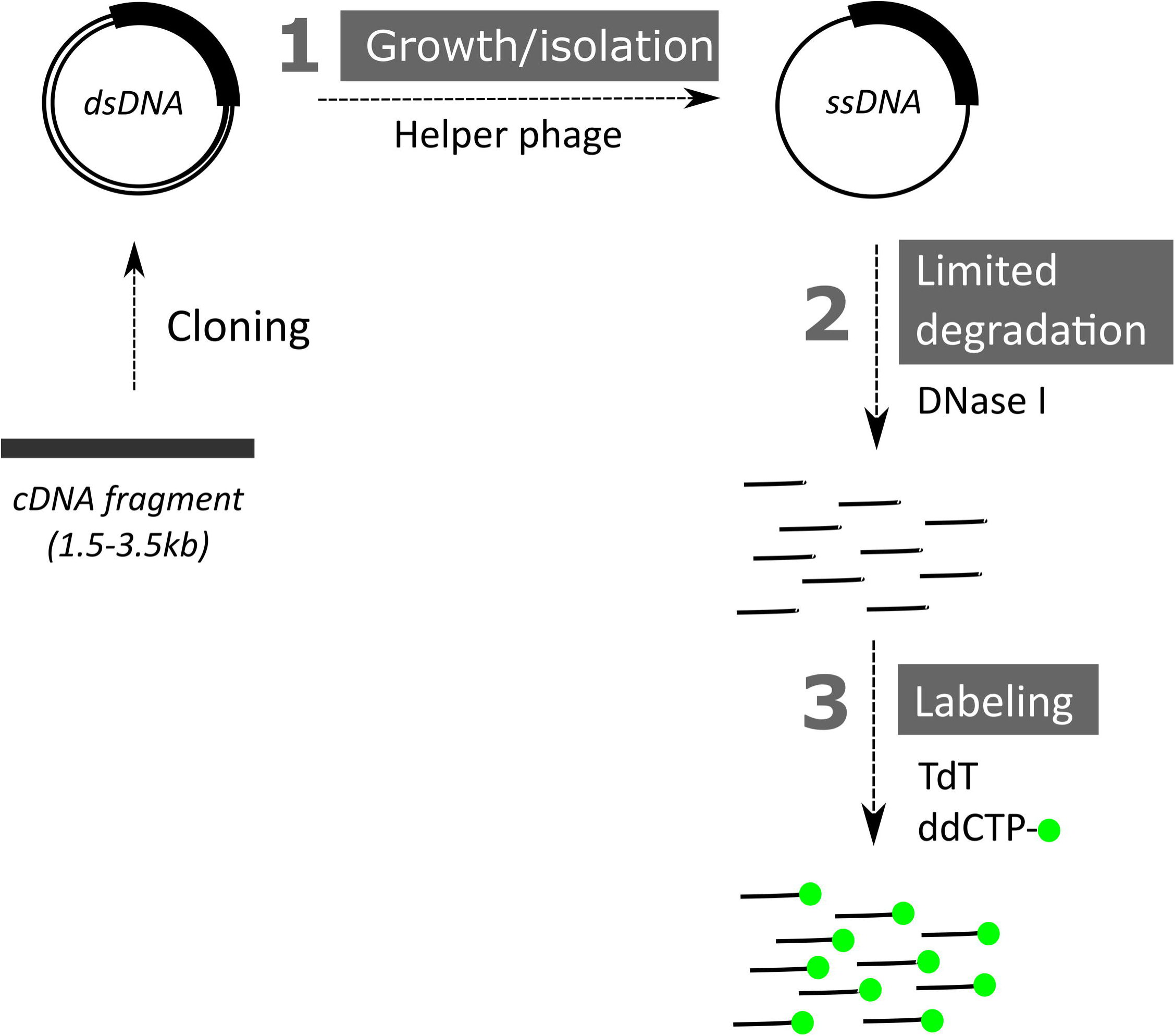
Scheme of enzyme-based synthesis of smRNA FISH probes. The method consists of the following main steps: **(1)** growing and isolating single stranded copies of a phagemid that comprises the target sequence; **(2)** fragmenting these single stranded DNA molecules; and **(3)** end-labeling the resulting oligonucleotides with a single fluorophore.

### Isolation of single stranded DNA molecules that comprise a segment that is complementary to the target RNA molecule

In smRNA FISH, the use of single-stranded probes ensures efficient hybridization to the target sequence. Since one of the goals in developing an enzyme-based method for the synthesis of smRNA FISH probes was to obviate the need to purchase pools of oligonucleotides, it was necessary to find a mean to generate single stranded DNA (ssDNA) that could be used as starting material. To that end, I decided to exploit the well-characterized properties of phagemids, which are vectors that can be replicated as double stranded molecules or, upon infection of cells with a helper phage, as single stranded ones [21]. These single stranded copies are packaged and exported in the culture medium, from which they can be easily recovered. Thus, the preliminary step in the protocol is to clone a fragment corresponding to the target RNA into a plasmid containing the origin of replication of the f1 or M13 bacteriophage, e.g. into the blunt SmaI site of the pGEM-3zf(-) phagemid from Promega. Vectors harboring the fragment in both orientations are easily obtained; if the fragment is cloned in the same orientation as the phage origin, the vector can be used to generate a control sense probe, if in the opposite orientation, an antisense probe. Since the whole fragment is used to derive probes (see below), care should be taken to avoid including elements that might hybridize to non-specific sequences, such as repeat regions, intron-exon boundaries, polyadenine stretches or long (> 50 bp) low complexity regions. Ideally, the cloned fragment should be between 1.5 and 3.5 kb in length and mostly correspond to the coding sequence of a mature mRNA transcript. Before cloning, the fragment sequence is always analyzed by BLAST against cDNA databases using stringent parameters; it is rejected if a given part of it is found to hybridize to multiple mRNAs, which can be the case in particular for conserved protein binding sites in the 3’ untranslated region. Once the phagemid has been engineered to harbor the probe fragment and transformed into an F factor-expressing *E. coli* strain (e.g. JM109), microgram quantities of ssDNA can be obtained and purified according to standard methods. I found that final purification of the recovered ssDNA molecules on commercially-available ion-exchange columns helped to remove contaminating traces of small DNA fragments or RNA which could complicate subsequent steps.

### Generation of short oligonucleotides from single stranded DNA molecules by limited DNase I digestion

In order to perform efficiently, the smRNA FISH probe must be made up of oligonucleotides that are typically between 20 and 60 nucleotides in length. The small size of the probes ensures their efficient penetration into the sample. Since the size of the ssDNA molecules isolated in the previous step is the size of the phagemid (3.2 kb) plus the size of the insert, it is necessary to fragment these molecules into oligonucleotides before proceeding to the labeling step. To do so, the ssDNA molecules are incubated with DNase I, a non-specific endonuclease that is active on single stranded DNA and leaves free 3’ hydroxyl groups on its products. Various conditions were tested to find the one that leads to fragmentation of the ssDNA molecules into oligonucleotides of approximately 20 to 60 nucleotides in length. It was found that incubating 1 μg of ssDNA with 50-100 mU (Kunitz) of DNase I in a total volume of 10 μl performed well. The reactions were assembled on ice and incubated for 15 minutes at 65°C. Under these conditions, the enzyme was minimally active and rapidly inactivated, thereby ensuring limited digestion of the ssDNA molecules and generation of fragments of the desired size. Figure 2 shows the result of a polyacrylamide gel analysis of ssDNA molecules comprising a 3.1 kb fragment complementary to the human large subunit of RNA polymerase II (hPOLR2A) open reading frame sequence and incubated with progressively higher amounts of DNase I. The pool for which the bulk size of oligonucleotides is between 20 and 60 nucleotides (lane 3 in this particular case) was chosen for labeling. Under these conditions, and assuming an average size of 35 bases, the amount of each oligonucleotide species present after fragmentation of 1 μg of the 6.2 kb ssDNA molecules (90 pmoles) is ~0.5 pmole and the total oligonucleotide concentration is ~ 9 μM.

With the present method, and assuming as above an average length of 35 bases, controlled digestion of ssDNA molecules can in theory give rise to up to 88 different oligonucleotides that cover the 3.1 kb fragment complementary to the target RNA. In practice, oligonucleotides generated by random cutting of a large fragment vary in size and overlap on the target sequence. Nonetheless, it is safe to assume that multiple (> 30) oligonucleotides derived from the ssDNA molecules can uniquely hybridize to the target sequence. Since their identity and their exact length remain unknown, these oligonucleotides will be referred to as ‘random’ oligonucleotides.

**Figure 2.**
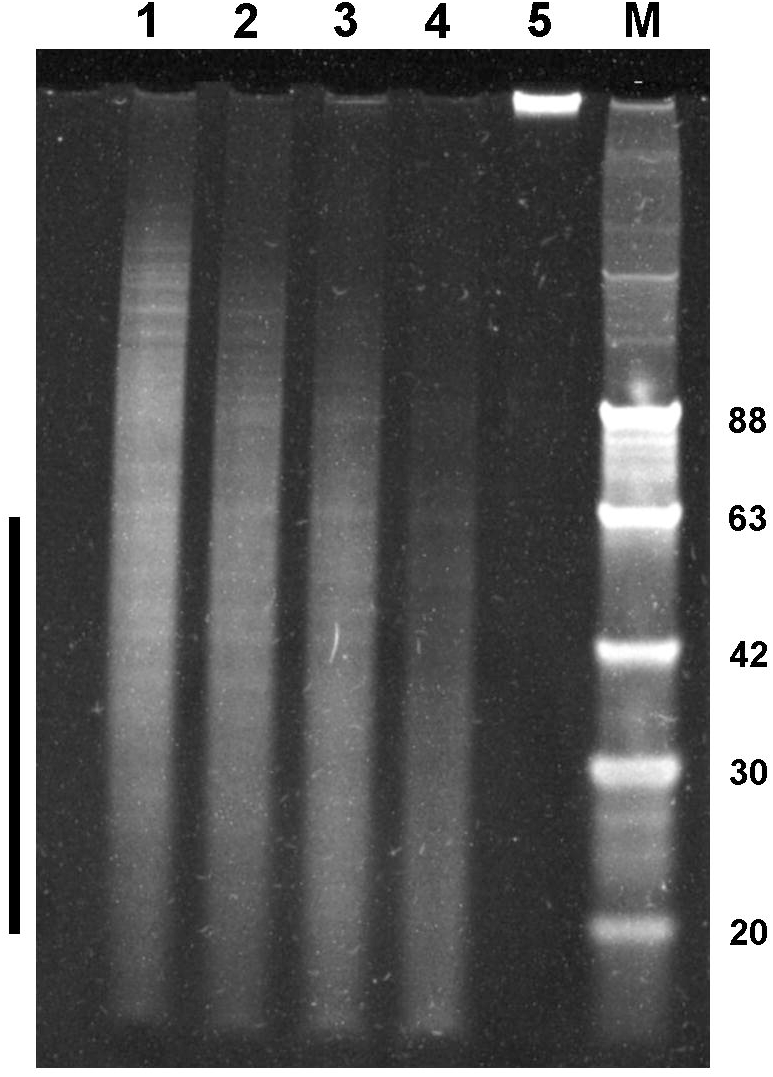
Limited digestion of single stranded DNA using DNase I. Shown is the length of fragments generated after incubation of single stranded DNA molecules with progressively higher concentration of DNase I. Reaction products (total of 100 ng) were separated on a denaturing 12% polyacrylamide gel. **Lane 1-4**: final DNase I concentrations of 2 mU/μl, 3.3 mU/μl, 5 mU/μl and 10 mU/μl. **Lane 5**: no DNase I. **Lane M**: equimolar mix of oligonucleotides of the indicated lengths. The ideal range of fragment lengths (~20-60 nucleotides) is indicated on the left.

### End labeling of oligonucleotides with fluorescent dideoxynucleotides

One of the breakthroughs in the development of the smRNA FISH technique was the realization that a probe consisting of multiple oligonucleotides labeled with a single fluorophore gave a more robust and reproducible signal than one consisting of only a few oligonucleotides labeled with multiple fluorophores [14]. The last step of the method presented here is therefore to label the oligonucleotides generated in the previous step in such a way that a single fluorophore is added per molecule. This is done using terminal deoxynucleotidyl transferase (TdT) and fluorescent dideoxynucleotides. The use of a dideoxynucleotide, which lacks a free 3’ hydroxyl group, ensures that one and only one label is added at the end of the oligonucleotides by the enzyme. Dye-labeled ddCTP, ddATP or ddGTP are preferred for the labeling since the affinity of TdT is higher for these analogs than it is for ddTTP [22]. Labeling is carried out by adding reagents directly to the inactivated DNase I reaction. Afterward, the oligonucleotides are separated from the bulk of unincorporated dye-labeled dideoxynucleotides by conventional techniques, e.g. ethanol precipitation or size exclusion chromatography. Measurement of the absorbance of a pool of Cy3-labeled oligonucleotides (average size estimated at 25 nt) at 260 nm and at 552 nm revealed that ~1.3 molecule of fluorophore was bound per oligonucleotide (Figure 3). This result indicates that TdT can be used to efficiently label DNA ends with a single fluorophore, in general agreement with published observations about this enzyme [23].

**Figures 3.**
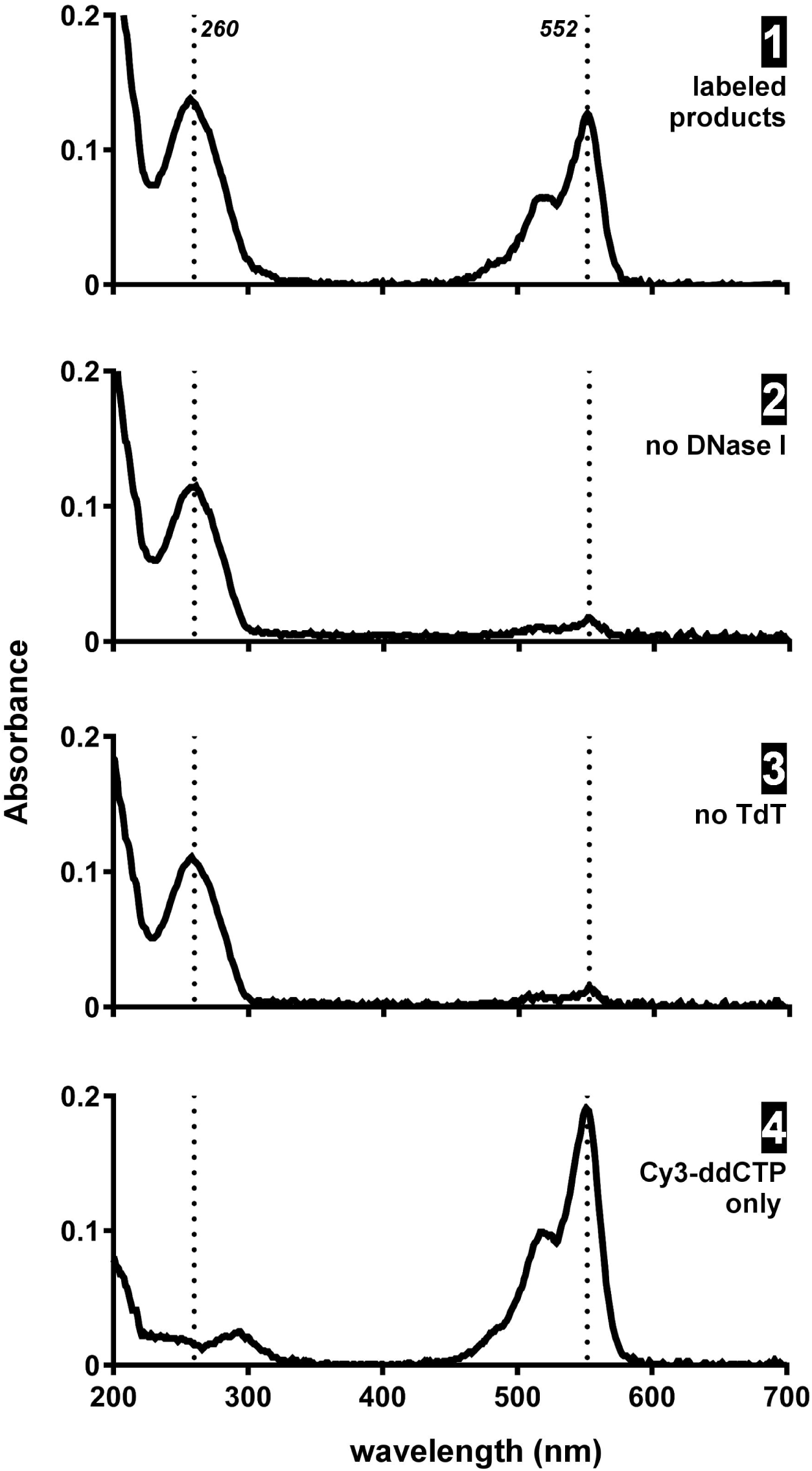
Absorption spectra of labeled and control reaction products. Labeling reactions with Cy3-ddCTP were performed on ssDNA molecules that underwent prior limited digestion (spectrum 1) or not (spectrum 2), in the presence (spectrum 1) or absence (spectrum 3) of TdT. The absorption spectrum of a 10 μM solution of Cy3-ddCTP is also shown (spectrum 4). The dotted lines indicate the absorption maxima for DNA (260nm) and Cy3 (552nm).

### Validation of the method

The first step in the validation of a new method is to see how it compares with established ones. To that end, HepG2 cells were co-labeled with two probes against the hPOLR2A transcript. One consisted of a pool of 48 oligonucleotides (20-mer) distributed over the entire 6.7 kb mRNA sequence, the other was generated by TdT-mediated labeling of DNase I-fragmented ssDNA molecules comprising a 3.1 kb fragment of hPOLR2A (nt 2091-5182) in the antisense or sense orientation. The first probe was labeled with Quasar-670 and is referred to as Stellaris, from its commercial name; the second was labeled with Cy3 and is referred to as TdT-labeled, from the main step of the method described here. Since the latter probe is made up of labeled oligonucleotides not only against the target sequence, but also against the phagemid sequence, an excess (~9-fold) of unlabeled competitor oligonucleotides was included in the hybridization mix. These were generated by controlled DNase I digestion of ssDNA molecules replicated from a phagemid without insert and with the f1 origin in the opposite orientation, i.e. pGEM3zf(+) in this case.

Figure 4A shows a typical smRNA FISH results obtained with the chemically-synthesized probe (Stellaris). Figure 4B shows the same field of view, but with smRNA FISH signals obtained with the TdT-labeled antisense probe. Both images show intense and distinct dots in the cytoplasm of the cells. The specificity of the TdT-labeled probe was ascertained by the lack of any dot-like signals from the sense probe (Figure 4C). Merging of the two positive images shows an excellent colocalization of signals obtained with the two different probes (Figure 4D-E). Image analysis of 4 image stacks containing a total of 3,947 individual dots labeled with the TdT-labeled probe revealed that 83 % ± 3 % of them colocalized with dots labeled with the Stellaris probe, a percentage comparable to the one previously obtained when assessing the performance of branched DNA smRNA FISH probes [17].

**Figure 4.**
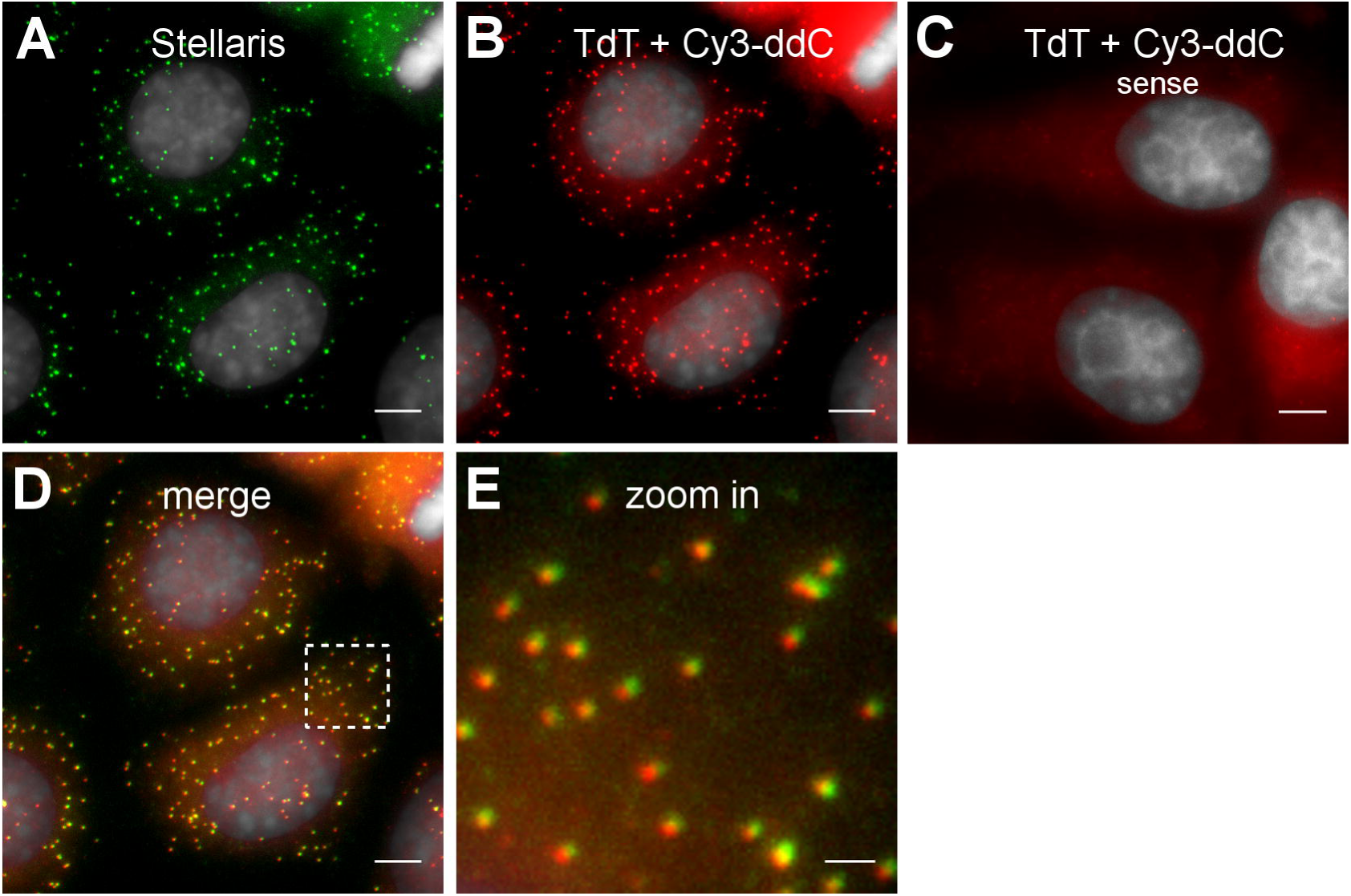
Comparison of smRNA FISH results obtained using commercially available
and enzyme-synthesized probes. HepG2 cells hybridized with two different probes against the human POLR2A mRNA. **(A)** Commercially available Quasar 670-labeled Stellaris probes of defined sequences. **(B)** Antisense TdT-labeled probes (Cy3 dye). **(C)** Sense TdT-labeled probes (Cy3 dye). **(D)** Merged images A+B. **(E)** Zoom in on the region boxed in D. Note the high level of co-localization of the two signals. Shown are maximum projections of optical sections. The cell nuclei were counterstained with DAPI (gray). Scale bar, 5 μm (A-D) or 1 μm (E).

To further characterize and compare the signals generated by the two types of probes, average mRNA signals were constructed using the MATLAB-based FISH-quant program [16]. Hybridization was carried out separately with single probes against POLR2A, both of which were labeled with near-infrared fluorophores (Cy5 and Quasar-670). The number of dots used in the averaging process was 4,322 for the Stellaris probe (Figure 5A-B). Measurement of the full-width at half maximum of intensity (FWHM) gave an average signal size of 354 nm (Figure 5C). The distribution of background-corrected signal amplitudes is shown in Figure 5D. In the case of the TdT-labeled probe, the number of dots used in the averaging process was 2,400 (Figure 5E-F). The average signal size was found to be 336 nm (Figure 5G), similar to the one obtained using the Stellaris probe. The distribution of background-corrected signal amplitudes for the TdT-labeled probe (Figure 5H) displays a peak at the same value as does the Stellaris probe. However, the distribution is more flattened for the TdT-labeled probe and the standard deviation is correspondingly greater (coefficient of variation of 0.41 vs. 0.25 for the Stellaris probe), consistent with a more heterogeneous composition of the TdT-labeled probe. Taken together, these results show that a probe obtained using the method described here performs as well as a chemically-synthesized probe obtained commercially.

**Figure 5.**
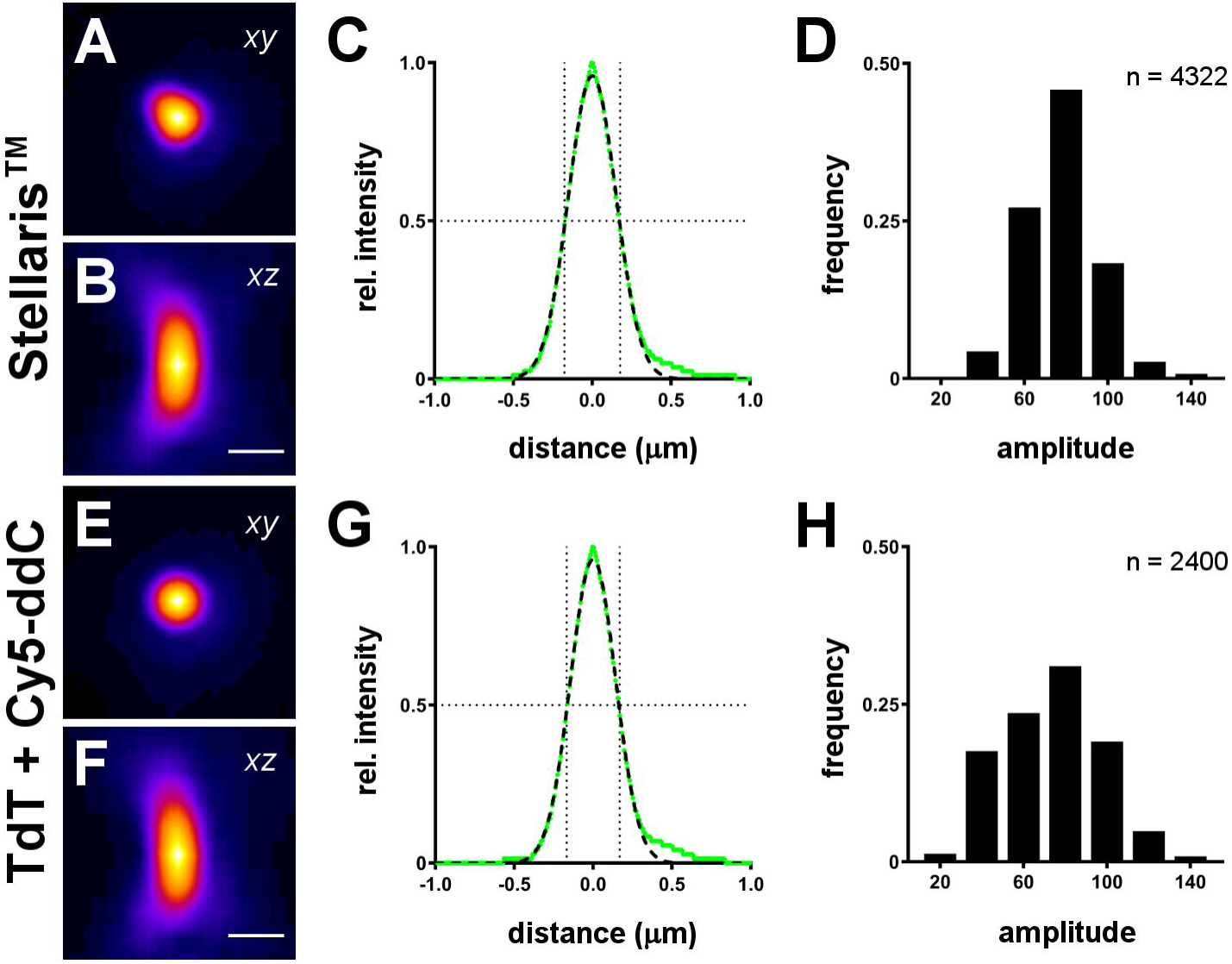
Size and amplitude distribution of smRNA FISH signals obtained using either commercially available or enzyme-synthesized probes. Size and amplitude distribution of smRNA FISH signals obtained using either commercially available or enzyme-synthesized probes. Average hPOLR2A mRNA signals were constructed from 4,322 or 2,400 dot-like signals obtained using a commercially available Stellaris probe **(A-B)** or a TdT-labeled probe **(E-F)**, respectively. Shown are *xy* and *xz* maximum projections. Scale bars, 0.5μm.**(C, G)** The relative intensity of these averaged signals was plotted (green lines) along a 2 μm *xy* line centered on the maximum value and the full width at half maximum (fine dotted lines) was measured on a Gaussian fit (thick dotted curves). **(D,H)** Distributions of background-corrected amplitudes of signals for each type of probes (bin size = 20). Note that the peak is in the same bin, but the variability is greater for the TdT-labeled probe.

To assess the specificity of smRNA FISH signals obtained using ‘random’ oligonucleotide probes labeled with TdT, I performed two-color labeling of the same hPOLR2A transcripts in HepG2 cells. As shown in Figure 6, probes labeled either with Cy5-ddCTP or with Cy3-ddCTP gave rise to the same signals, albeit with more or less inversely correlated intensities, presumably because of competition between the probes for binding to the same transcript. Image analysis of 5 image stacks containing a total of 3,929 individual Cy3-labeled dots revealed that 94 % ± 2 % of them colocalized with Cy5-labeled dots. The co-labeling of the same RNA molecules with probes that were independently synthesized and that harbor spectrally distinct dyes is evidence for the high specificity of the enzyme-synthesized smRNA FISH probes for their target.

**Figure 6.**
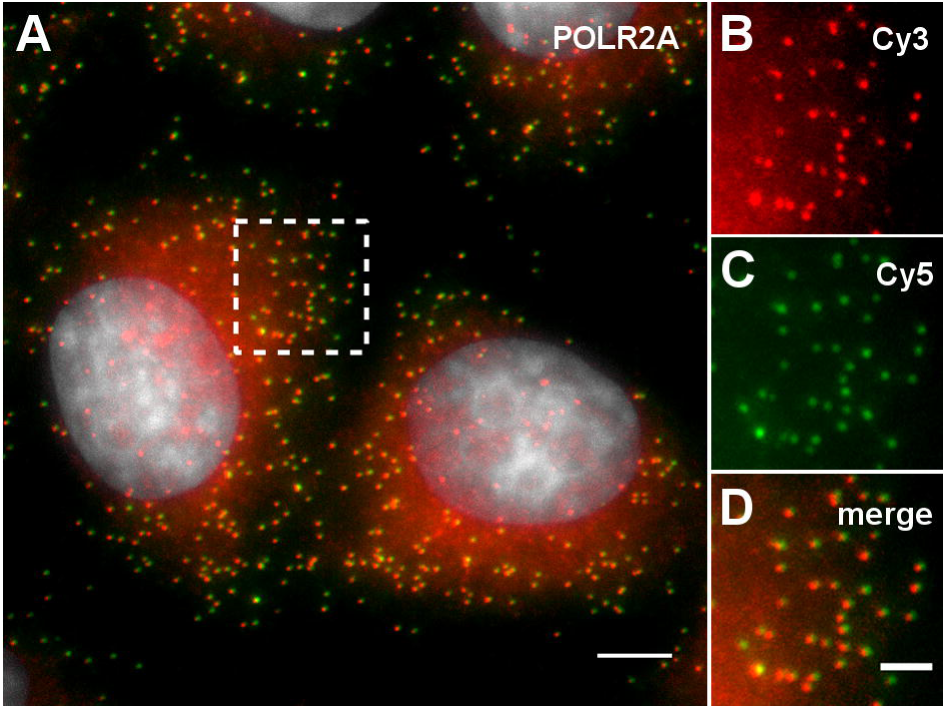
Specificity of enzyme-synthesized smRNA FISH probes. HepG2 cells were labeled simultaneously with Cy3-labeled (red) and Cy5-labeled (green) probes against POLR2A transcripts. Shown are maximum projections of optical sections. **(A)** Merged signals from both probes. The cell nuclei were counterstained with DAPI (gray). Scale bar, 5 μm. **(B-D)** Zoom in on the region boxed in A. The individual signals are shown, as well as the merged image. Scale bar, 2 μm.

One of the strengths of the smRNA FISH technique is the ability to detect multiple RNA species in the same cell. It was therefore of interest to generate pools of random oligonucleotides against different transcripts and to label them with spectrally distinct fluorophores. The fragments that were used as starting material in the generation of these probes varied in size from 1.4 kb (CREB3) to 3 kb (POLR2A), but their proportion of G+C bases were similar (56-58%). All probes gave nice dot-like signals under standard hybridization and washing conditions (Figure 7). As expected, the signal-to-background ratio was proportional to the length of the fragment cloned in the phagemid vector. Although the ratio was on average only around 1.1 for the smallest fragment (CREB3), the dots could nonetheless be easily recognized and counted by the FISH-QUANT smFISH algorithm. The rapid detection of multiple transcripts without the need for any optimization step in probe synthesis or hybridization shows that the technique of smRNA FISH probe synthesis described here is versatile and efficient.

**Figure 7.**
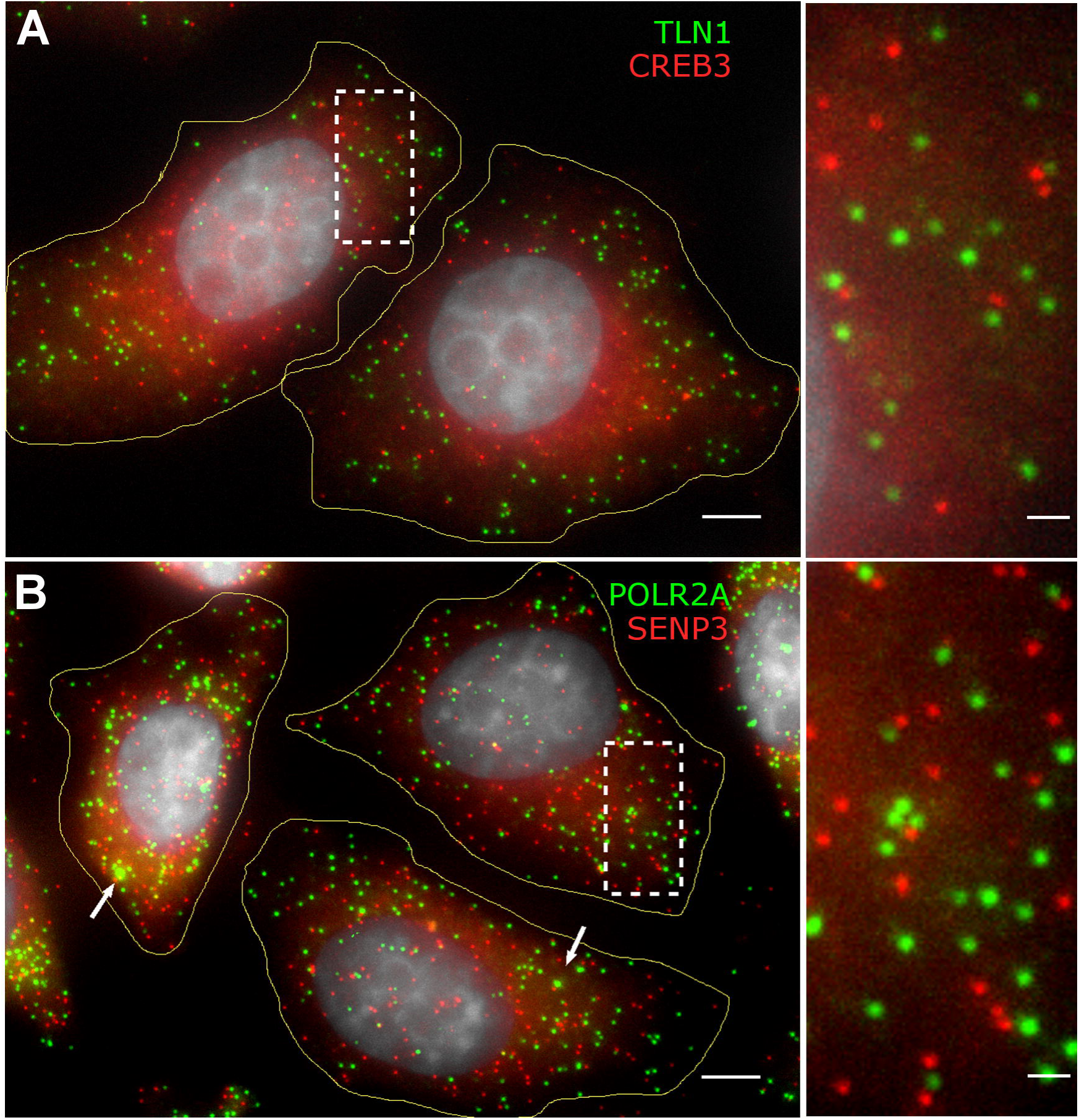
Detection of multiple transcripts in HepG2 cells. Probes targeting fragments of TLN1 and CREB3 transcripts **(A)** or POLR2A and SENP3 transcripts **(B)** were TdT-labeled with Cy5 (green) or Cy3 (red). Shown are maximum projections of optical sections. The cell nuclei were counterstained with DAPI (gray) and the cell contours are outlined. Scale bar, 5 μm. The panels on the right show digital enlargments of the boxed regions. Scale bar, 1 μm.

## DISCUSSION

This paper describes a novel method to generate probes for smRNA FISH. It uses standard molecular biology techniques and obviates the need to purchase pools of labeled oligonucleotides. It can be argued that nothing can be simpler nor more efficient than to order a ready-to-use smRNA FISH probe from a commercial source. However, I have found that the high price of these reagents warranted the investment needed to synthesize smRNA FISH probes using the technique described here. Once implemented, in house enzyme-based synthesis of probes has proven to be rapid and cost-effective. Multiple smRNA FISH probes can be generated in parallel in less than a week and, while it is true that fluorescent dideoxynucleotides are expensive reagents, the amount needed per reaction cost less than 25 euros. An additional advantage of the technique is the flexibility it affords to label the same fragmented ssDNA molecules with different dyes. Furthermore, the fact that a sense probe can be easily generated alongside its antisense counterpart can also prove useful, especially in the case of weakly expressed genes for which negative controls might be needed.

The technique consists of 3 main steps, each of which deserves some technical comments. The aim of the first step is to generate ssDNA molecules that comprise a fragment complementary to the target RNA. This is achieved through infection of phagemid-containing cells with a helper phage. Some of the problems that are often encountered, e.g. low yield of ssDNA and contamination with genomic and/or helper phage DNA, were avoided by early infection of the cell culture and growth at a lower temperature [18, 19]. The second step is the limited digestion of the ssDNA with DNase I to generate a pool of oligonucleotides ~20-60 nucleotides in length. Obviously, the optimal digestion conditions have to be determined empirically, but I found that, once they have been, the reaction is quite reproducible. To be on the safe side, I always perform two reactions with 2 different dilutions of DNase I so that at least one of them contains a pool of oligonucleotides of the desired length. The third and last step is the end-labeling of the oligonucleotides with a single fluorescent moiety. This is accomplished using TdT and dye-labeled dideoxynucleotide. At the concentrations of substrate that are used (100 μM for the labeled ddCTP and ~9 μM for the oligonucleotides), the reaction proceeds extremely efficiently [22].

It is important to stress that, contrary to chemically-synthesized oligonucleotide probes which have defined lengths and sequences, the ones that are generated using DNase I and TdT are of varying lengths and sequences (and hence G+C contents). In theory, the hybridization conditions, i.e. mainly the concentration of formamide, can be adjusted to obtain the best possible signal-to-background ratio. In practice, however, I found that this parameter did not need to be optimized provided that the bulk of oligonucleotides was smaller than 80 bases. The reason for this lies in the nature of the smRNA FISH signals, which arise only if a sufficient number of labeled oligonucleotides hybridize to the same mRNA molecule [14]. Thus, targeted hybridization events are averaged while non-specific ones do not give rise to signals above background because they do not occur on the same molecule. The variable length of the TdT-labeled oligonucleotides results in a greater variability in signal intensity, but not in a higher proportion of false-positive signals, as was found in the case of POLR2A (Figures 4 and 5).

The fact that the sequences of smRNA FISH probes generated through random DNase I digestion of ssDNA molecules are not defined deserves more attention. On the one hand, the high complexity of the TdT-labeled probes can turn out to be an advantage. Indeed, since the efficiency of hybridization depends in part on the secondary structure in the target RNA, it can be assumed that a pool of ‚random‘ oligonucleotides is more likely to cover a greater fraction of the target mRNA than is a pool of known sequences. On the other hand, care should be taken to avoid including low-complexity sequences in the fragment chosen to be complementary to the target RNA. So far, I have tested more than 15 different smRNA FISH probes that were generated using the present method. Only one gave unsatisfactorily high background, presumably due to the presence of long stretches of Ts which hybridized with cellular polyA tails. Another factor that is determinant in the performance of TdT-labeled smRNA FISH probes is the length of the fragment that is cloned into the phagemid and that is complementary to the target RNA. After limited digestion with DNase I, a 1-kb fragment can in theory give rise to 33 different oligonucleotides with an average length of 30 nucleotides while a 2-kb fragment can give rise to twice that number. The smallest fragment that was tested was 1.4 kb in length (to detect human CREB3 transcripts); it gave satisfactory results (Figure 7). The mean size of human mRNAs is 2.3 kb [24]. It is therefore possible to design useful probes against most mRNAs. However, when atttempting to detect smaller RNA targets, such as non coding RNAs, or intronic sequences, for which it is often difficult to find a long fragment without any repeats, the fact that the oligonucleotide probes will be derived from a smaller fragment means a lower signal strength. In this case, it may be useful to isolate oligonucleotides of a defined size (e.g. 20-mer) from the pool of labeled ones in order to limit the competition from longer oligonucleotides during hybridization and thus maximize the number of fluorophores per single target RNA molecule.

In summary, I have described a novel method to generate smRNA FISH probes using standard molecular biology techniques. The method is straightforward, simple and cost-effective. Its performance is comparable to that of chemically-synthesized dye-labeled oliogonucleotide pools. Single molecule RNA FISH is an imaging technique that provides quantitative information on gene expression at the level of single cells in their native environment. As such, it has provided important insights into the processes of gene regulation. It is hoped that the method of probe synthesis presented here will allow even more researchers to benefit from the use of the powerful smRNA FISH technique.

## ACKNOWLEDGEMENTS

This work was supported by the Ministry of Education, Youth and Sports of the Czech Republic within the LQ1604 National Sustainability Program II (Project BIOCEV-FAR) and by the project BIOCEV (CZ.1.05/1.1.00/02.0109).

---

**SUPPL TABLE 1.**
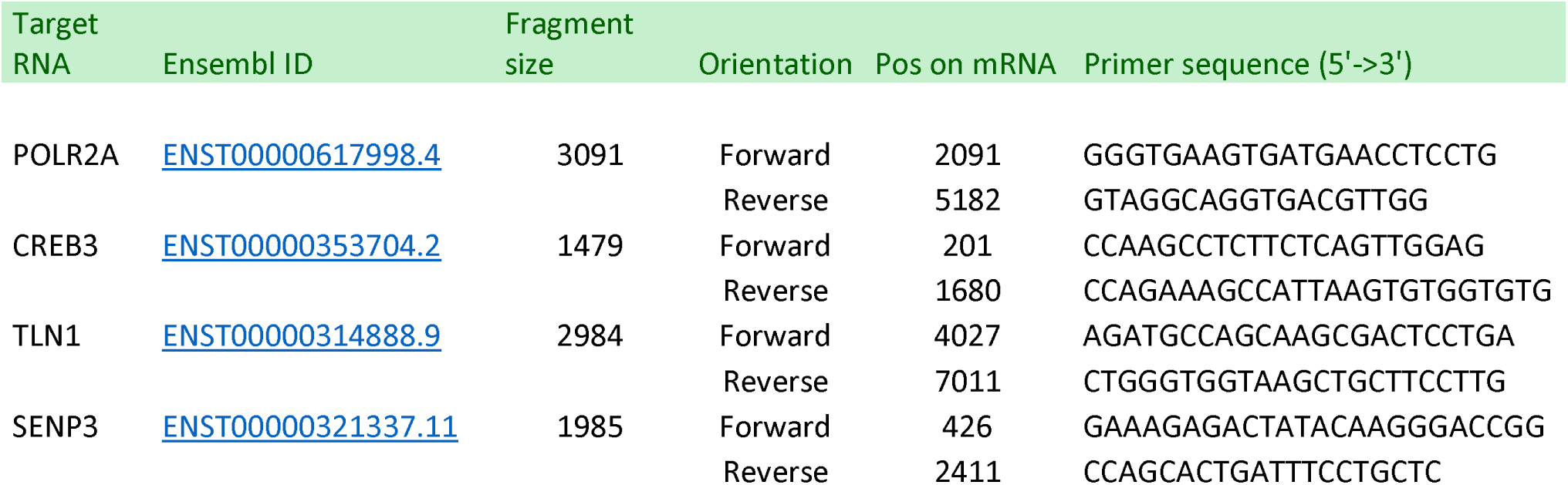
PRIMERS USED TO AMPLIFY FRAGMENTS COMPLEMENTARY TO TARGET RNAs.

